# Rbm45 Phylogenetics, Protein Domain Conservation, and Gene Architecture in Clade Metazoa

**DOI:** 10.1101/2024.02.29.582754

**Authors:** Virdjinija Vuchkovska, Teagan N. Marti, Anali Cisneros, Lauren M. Saiki, Jeffrey O. Henderson

**Author notes:** Equal contribution (Virdjinija Vuchkovska, Teagan N. Marti, Anali Cisneros, Jeffrey O. Henderson).

## Abstract

Mammalian Rbm45 is predominately expressed in neuronal tissue and is integral in brain development and neuronal differentiation under physiological conditions. Dysregulation of Rbm45 has been strongly associated with neurodegenerative disorders in humans and can drive hepatocellular carcinoma through reprogramming lipid metabolism. Intriguingly, Rbm45 is an ancient protein, evolutionarily conserved throughout metazoans, including in sponges which lack a nervous system. Curiously, the evolution of Rbm45 gene structure and protein domain conservation across kingdom Animalia is largely unknown. We performed phylogenetic analysis of Rbm45 nucleotide and amino acid sequences from 36 species representing 9 phyla: Porifera, Cnidaria, Priapulida, Mollusca, Brachiopoda, Arthropoda, Echinodermata, Hemichordata, and Chordata. While the tree from Rbm45 nucleotide sequence data resulted in clades Protostomia and Deuterostomia showing paraphyly, the phylogeny derived from Rbm45 amino acid sequence largely recapitulated known monophyletic relationships among metazoans. Human RBM45 protein structure includes three RNA-binding domains (RBD), a homo-oligomerization association (HOA) domain, a nuclear localization sequence (NLS), and a nuclear export sequence (NES). Multiple sequence alignment across the same 36 taxa used for phylogenetic analysis revealed conservation of all three RBDs, the HOA, and NLS; in contrast the NES was only detected in clade Craniata and not in clades Ambulacraria and Protostomia. *Rbm45* gene structure analysis revealed increasing gene complexity concomitant with increasing evolutionary complexity. *Rbm45* from non-bilaterian taxa had from 2 to 4 large exons, while bilaterian taxa had between 6 to17 small exons. These findings demonstrate that *Rbm45* is an ancient, highly conserved gene among metazoans suggesting a function in a breadth of neural/sensory systems.

## Introduction

RNA-binding proteins (RBP) are an evolutionarily conserved [1] family of proteins that have been shown to participate in a constellation of cellular functions. Specifically, RNA recognition motif-type (RRM) binding domain proteins (RBDPs) have been demonstrated to regulate post-transcriptional RNA metabolism by modulating the longevity [2–5] and translational efficacy [6–11] of target mRNAs and contributing to splicing reactions [12]. RBDPs have also been identified participating in RNA-mediated [13–18] and RNA-independent protein-protein complexes [19–21]. These biochemical properties of RBDPs have been linked to roles in apoptosis [22], tumorigenesis [23], and neuropathology [24,25].

*Rbm45* is an ancient gene, conserved from sponges to humans [26]. The human *RBM45* locus on chromosome 2q31.2 [26] encodes a promiscuous RBDP found to be expressed under normal physiological conditions predominantly in neuronal tissue of rats [27], mice [26,28], and humans [29,30]. Rbm45 has three canonical RNA-binding domains (RBD I, II, III) [27,31,32] that preferentially bind GC-rich RNA sequence motifs [27,33,34] and have recently been shown to facilitate binding to single-stranded DNA [35]; additionally, N-terminal RBD I and RBD II have been demonstrated to participate in RNA-independent protein-protein interactions [36].

Furthermore, Rbm45 contains a homo-oligomer assembly (HOA) domain that mediates homo- and heteromerization of Rbm45 and binding partners [31,37]. Finally, Rbm45 is capable of shuttling between the nucleus and cytoplasm by means of a nuclear localization signal (NLS) [31,32] and a nuclear export signal (NES) [32] facilitating its presumed role in regulating nuclear spliceosome and mRNA splicing functions [37–40].

Rbm45 was initially identified as being expressed spatiotemporally in neuronal precursor cells during brain development in rats [27] and has recently been shown to be necessary for neuroblastoma differentiation in the SH-SY5Y cell line [41]. Early work associated upregulation of Rbm45 with suppression of hypoxia induced apoptosis of cardiomyocytes [42] in mice and neuronal repair during spinal cord injury in a neonatal opossum model [43]. Subsequently, biochemical [25,29,31,32,37,44] and molecular genetic [45] studies have linked human RBM45 protein dysfunction/aggregation to the neurodegenerative syndromes: frontotemporal dementia, amyotrophic lateral sclerosis, and Alzheimer’s disease. These data implicate Rbm45 in neurogenesis, neural repair, and neuropathophysiology. We [26] and others [31,32] have shown that *Rbm45* is highly conserved across metazoan taxa; intriguingly, our previous work [26] identified an *Rbm45* orthologue in sponges (phylum Porifera) which lack a nervous system.

However, sponges do have a neural toolkit of genes [46] as well as neuroid cells [47] with secretory vesicles capable of communicating with adjacent cell types and coordinating cellular activity; this is suggestive of a vital role for phylum Porifera in nervous system evolution. To explore the deep homology of *Rbm45* in neural development, we have analyzed the phylogenetic history, protein domain conservation, and gene structure across 36 Rbm45 orthologues from 9 metazoan phyla. In this study, we show conservation of *Rbm45* over 650 million years of evolutionary history with preservation of RNA-binding, HOA, and NLS regions, as well as increasing complexity of gene architecture during the radiation of clade Metazoa.

## Experimental Procedures

### Rbm45 Nucleic Acid and Amino Acid Sequences

All Rbm45 sequence information (genomic; cDNA; amino acid) in this study was retrieved from the National Center for Biotechnology Information (NCBI; https://www.ncbi.nlm.nih.gov/). *Rbm45* orthologues [48] were identified using the BLAST algorithm by querying GenBank with the *Homo sapiens* (human; Gene ID:129831) *RBM45* cDNA, accession number (no.) NM_152945.3 [26]. The following accession nos. were utilized in this study: phylum Porifera: *Amphimedon queenslandica* (sponge: NW_003546244.1; XM_003382583.3; XP_003382631.1); phylum Brachiopoda: *Lingula anatine* (lamp shell: NW_019775657.1; XM_013560776.1; XP_013416230.1); phylum Priapulida: *Priapulus caudatus* (priapulid worm: NW_014578398.1; XM_014820871.1; XP_014676357.1); phylum Hemichordata: *Saccoglossus kowalevskii* (acorn worm: NW_003156738.1; XM_006825506.1; XP_006825569.1); phylum Echinodermata: *Acanthaster planci* (crown-of-thorns starfish: NW_019091356.1; XM_022231661.1; XP_022087353.1), *Strongylocentrotus purpuratus* (purple sea urchin: NW_011995289.1; XM_780088.4; XP_785181.2); phylum Cnidaria: *Exaiptasia pallida* (pale anemone: NW_018384422.1; XM_021061761.1; XP_020917420.1), *Hydra vulgaris* (freshwater polyp: NW_004169798.1; XM_012704848.1; XP_012560302.1), *Stylophora pistillata* (smooth cauliflower coral: NW_019217956.1; XM_022936599.1; XP_022792334.1), *Orbicella faveolata* (mountainous star coral: NW_018149652.1; XM_020763593.1; XP_020619252.1), *Acropora digitfera* (staghorn coral: NW_015441794.1; XM_015923864.1; XP_015779350.1); phylum Mollusca: *Octopus bimaculoides* (California two-spot octopus: NW_014653530.1; XM_014915327.1; XP_014770813.1), *Aplysia californica* (California sea hare: NW_004797628.1; XM_013085237.1; XP_012940691.1), *Crassostrea gigas* (Pacific oyster: NW_011936719.1; XM_011449747.2; XP_011448049.1); phylum Arthropoda: *Limulus polyphemus* (Atlantic horseshoe crab: NW_013677256.1; XM_013937428.2; XP_013792882.2), *Bactrocera dorsalis* (oriental fruit fly: NW_011876306.1; XM_011202560.2; XP_011200862.1), *Drosophila melanogaster* (fruit fly: NT_037436.4; NM_139630.3; NP_647887.1), *Aedes aegypti* (yellow fever mosquito: NC_035108.1; XM_001660954.2; XP_001661004.2); phylum Chordata: *Branchiostoma floridae* (Florida lancelet: NW_003101314.1; XM_002586844.1; XP_002586890.1), *Latimeria chalumnae* (West Indian Ocean coelacanth: NW_005822196.1; XM_006012287.1; XP_006012349.1), *Callorhinchus milii* (Australian ghost shark: NW_006890067.1; XM_007890069.1; XP_007888260.1), *Rhincodon typus* (whale shark: NW_018067517.1; XM_020531702.1; XP_020387291.1), *Orcinus orca* (killer whale: NW_004438435.1; XM_004267367.2; XP_004267415.1), *Lipotes vexillifer* (Yangtze River dolphin: NW_006776904.1; XM_007450717.1; XP_007450779.1), *Chelonia mydas* (green sea turtle: NW_006631183.1; XM_027822939.1; XP_027678740.1), *Alligator mississippiensis* (American alligator: NW_017714267.1; XM_019480600.1; XP_019336145.1), *Xenopus laevis* (African clawed frog: NC_030741.1; NM_001086621.1; NP_001080090.1), *Danio rerio* (Zebrafish: NC_007120.7; NM_001127402.1; NP_001120874.1), *Ornithorhynchus anatinus* (platypus: NW_001794453.1; XM_016228192.1; XP_016083678.1), *Monodelphis domestica* (gray short-tailed opossum: NC_008804.1; XM_007494431.2; XP_007494493.1), *Canis lupus familiaris* (dog: NC_006618.3; XM_022414739.1; XP_022270447.1), *Rattus norvegicus* (Norway rat: NC_005102.4; NM_153306.1; NP_695218.1), *Mus musculus* (house mouse: NC_000068.7; NM_153405.2; NP_700454.1), *Gallus gallus* (chicken: NC_006094.5; NM_001031252.1; NP_001026423.1), *Loxodonta africana* (African savanna elephant: NW_003573423.1; XM_023543049.1; XP_023398817.1), *Gorilla gorilla* (western gorilla: NC_018426.2; XM_019021614.1; XP_018877159.1), *Pan troglodytes* (common chimpanzee: NC_036881.1; XM_515938.6; XP_515938.2), *Homo sapiens* (human: NC_000002.12; NM_001365578.1; NP_001352507.1).

### Molecular Phylogenetic Trees

All molecular phylogenetic trees were created using Molecular Evolutionary Genetic Analysis v.7.0 software (MEGA7) [49]. cDNA sequence alignments were performed using ClustalW within the MEGA7 program using these default parameters: Pairwise Alignment-Gap Opening Penalty: 15, Gap Extension Penalty: 6.66; Multiple Alignment-Gap Opening Penalty: 15, Gap Extension Penalty: 6.66; DNA Weight Matrix: IUB; Transition Weight: 0.5; Use Negative Matrix: OFF; Delay Divergent Cutoff (%): 30. Molecular phylogenetic tree creation with Rbm45 cDNA sequences used the following parameters: Statistical Method: Maximum Likelihood [50]; Test of Phylogeny: Bootstrap method [51], No. of Bootstrap Replications: 100 and 1000; Model: Tamura-Nei model; Rate among Sites: Uniform rates; Gaps/Missing Data Treatment: Use all sites; ML Heuristic Method: Nearest-Neighbor-Interchange; Initial Tree for ML: NJ/BioNJ; Codons Included: 1^st^+2^nd^+3^rd^+Non-Coding.

Amino acid sequence alignments were executed using ClustalW within the MEGA7 program and used the following default parameters: Pairwise Alignment-Gap Opening Penalty: 10, Gap Extension Penalty: 0.1; Multiple Alignment-Gap Opening Penalty: 10, Gap Extension Penalty: 0.2; Protein Weight Matrix: Gonnet; Residue-specific Penalties: ON; Hydrophilic Penalties: ON; Gap Separation Distance: 4; End Gap Separation: OFF; Genetic Code Table: Standard; Use Negative Matrix: OFF; Delay Divergent Cutoff (%): 30. Molecular phylogenetic tree creation with Rbm45 amino acid sequences used the following parameters: Statistical Method: Maximum Likelihood [52]; Test of Phylogeny: Bootstrap method [51], No. of Bootstrap Replications: 100 and 1000; Model: Jones-Taylor-Thornton (JTT) model; Rate among sites: Uniform rates; Gaps/Missing Data Treatment: Use all sites; ML Heuristic Method: Nearest-Neighbor-Interchange; Initial Tree for ML: NJ/BioNJ.

### Rbm45 Orthologue Protein Domain Conservation

Multiple sequence alignments were performed using Clustal Omega (https://www.ebi.ac.uk/Tools/msa/clustalo/) [53–55] with the following parameters: Output Format: ClustalW with character counts; Dealign input sequences: No; Mbed-like clustering guide: Yes; Mbed-like clustering: Yes; Number of combined: default (0); Max guide tree iterations: Default; Max HMM Iteration: Default; Order: input.

NCBI gene annotation (https://www.ncbi.nlm.nih.gov/homologene/?term=Rbm45 [accessed 2024 February 4]), our previous work [26], and data from Tamada *et al.* [27] reveal that Rbm45 contains four RBDs. However, ensuing work by two leading Rbm45 research teams show an alternative structure with three RBDs [31,32]; for consistency, we have adopted these research team’s Rbm45 domain nomenclature for this study. RBDs I, II, and III, HOA domain, NES, and the monopartite NLS of Rbm45 were identified by visual inspection after performing multiple sequence alignment using human RBM45 RBDs, HOA, NES, and NLS sequences as reference [21,26,32]. Additionally, the Locating Nuclear Export Signals (LocNES) algorithm was used to extend the analysis of NES sequences [56]. Orthologous *Rbm45* exon and intron sequence lengths were retrieved from NCBI Genome (https://www.ncbi.nlm.nih.gov/genome/).

### Rbm45 Orthologue Gene Architecture Evolution

The Mean Exon Size versus Exon Number and Age of Taxonomic Lineage versus Exon Number were calculated for the following representative animals across metazoan taxa (common name; exon number; mean exon size in base pairs; approximate lineage age in years): phylum Porifera: *Amphimedon queenslandica* (sponge; 2; 984; 650,000,000 [57]); phylum Brachiopoda: *Lingula anatine* (lamp shell; 11; 238; 66,000,000 [58]); phylum Hemichordata: *Saccoglossus kowalevskii* (acorn worm; 9; 234; 373,000,000 [59]); phylum Echinodermata: *Acanthaster planci* (crown-of-thorns starfish; 11; 404; 4,000,000 [60]), *Strongylocentrotus purpuratus* (purple sea urchin; 10; 183; 199,000,000 [61]); phylum Cnidaria: *Exaiptasia pallida* (pale anemone; 4; 544; 500,000,000 [62]), *Hydra vulgaris* (freshwater polyp; 3; 420; 540,000,000 [63]), *Orbicella faveolata* (mountainous star coral; 3; 937; 570,000,000 [64]); phylum Mollusca: *Octopus bimaculoides* (California two-spot octopus; 10; 219; 155,000,000 [65]), *Crassostrea gigas* (Pacific oyster; 13; 195; 15,000,000 [66]); phylum Arthropoda: *Limulus polyphemus* (Atlantic horseshoe crab; 6; 206; 250,000,000 [67]), *Drosophila melanogaster* (fruit fly; 9; 174; 5,400,000 [68]); phylum Chordata: *Branchiostoma floridae* (Florida lancelet; 7; 154; 100,000,000 [69]), *Latimeria chalumnae* (West Indian Ocean coelacanth; 10; 181; 70,000,000 [70]), *Orcinus orca* (killer whale; 10; 182; 11,000,000 [71]), *Lipotes vexillifer* (Yangtze River dolphin; 10; 170; 11,000,000 [71]), *Alligator mississippiensis* (American alligator; 10; 210; 53,000,000 [72]), *Xenopus laevis* (African clawed frog; 11; 173; 18,000,000 [73]), *Danio rerio* (Zebrafish; 10; 199; 150,000,000 [74]), *Monodelphis domestica* (gray short-tailed opossum; 10; 196; 3,000,000 [75]), *Rattus norvegicus* (Norway rat; 10; 178; 2,000,000 [76]), *Mus musculus* (house mouse; 10; 190; 6,000,000 [77]), *Gallus gallus* (chicken; 10; 180; 21,000,000 [78]), *Pan troglodytes* (common chimpanzee; 10; 181; 8,000,000 [79]), *Homo sapiens* (human; 10; 149; 300,000 [80]). Scatter plots and regression analysis (R^2^: Coefficient of Determination) were generated in Microsoft Excel.

## Results and Discussion

### Molecular Phylogenetics of Rbm45 Orthologues

Our lab [26] has previously demonstrated that nucleotide and amino acid phylograms of 10 vertebrate *Rbm45* orthologues recapitulate accepted taxonomic relationships between classes Actinopterygii (ray-finned fishes), Amphibia, Reptilia, and Mammalia; additionally, through NCBI database interrogation, we reported *Rbm45* orthologues, both empirically confirmed and Gnomon algorithm predicted (https://www.ncbi.nlm.nih.gov/genome/annotation_euk/process/), across metazoan taxa including animals from the non-Bilateria phyla Porifera (sponges) and Cnidaria (e.g., hydra), and within clade Bilateria phyla from the nephrozoan lineages Protostomia (e.g., phyla Mollusca and Arthropoda) and Deuterostomia (e.g., phyla Echinodermata and Chordata). Molecular phylogenetic analysis has been successfully used to reconstruct the evolutionary history of populations, genes, and proteins; furthermore, it has been utilized to understand genome organization and gene conservation [81,82]. Therefore, to gain a better understanding of the evolutionary history of *Rbm45*, a gene involved in neuronal development [27] and neuronal pathogenesis [29], we have expanded our phylogenetic analysis to 36 *Rbm45* orthologues from 9 phyla: Porifera, Cnidaria, Brachiopoda, Mollusca, Arthropoda, Echinodermata, Priapulid, Hemichordata, and Chordata. When more than one organism’s *Rbm45* sequence was available within a phylum, we often chose those species that are on the IUCN Red List of Threatened Species (e.g., *Lipotes vexillifer* [Yangtze River dolphin] [83], *Loxodonta africana* [African savanna elephant] [84]), are molecular genetic and developmental biology model systems (e.g., *Danio rerio* [Zebrafish], *Xenopus laevis* [African clawed frog], *Gallus gallus* [chicken], and *Mus musculus* [house mouse]), or give us multiple subgroups in a taxonomic unit (e.g., class Mammalia: order Monotremata [*Ornithorhynchus anatinus*: platypus], infraclass Marsupialia [*Monodelphis domestica*: gray short-tailed opossum], and infraclass Placentalia [*Homo sapiens*: humans]). Additionally, we attempted to have representative members from a variety of crown clades [85]: Ecdysozoa (phyla Priapulida and Arthropoda), Spiralia (phyla Brachiopoda and Mollusca), Ambulacraria (phyla Echinodermata and Hemichordata), and Chordata (subphylum Craniata) [86,87].

We used MEGA7 software (Materials and Methods) to build rooted phylogenetic trees using 36 Rbm45 cDNA and amino acid orthologous sequences. The bootstrap consensus tree from 1000 iterations is taken to represent the evolutionary history of the gene [51]. In our unbiased tree analysis (data not shown), phylum Porifera resolved as the sister group to all other animals; therefore, phylum Porifera served as the outgroup [88–90] in the cDNA and amino acid molecular phylogenies (Figs. 1 and 2). As we progress “up” the tree from most ancient to most recent lineages, phylum Cnidaria exhibits paraphyly, with the freshwater polyp (*Hydra vulgaris*) diverging after corals and anemones as sister group to Bilateria, having 87% and 97% bootstrap support for the evolutionary node in the cDNA and amino acid molecular phylogenies, respectively (Figs. 1 and 2). These data are an example of incomplete lineage sorting, where a gene tree does not match the history of the taxa [82,91], within clade Cnidaria [92]. Where we have sequence from more than one organism in a phylum, the phyla are monophyletic in both the cDNA and amino acid molecular phylogeny (e.g., Mollusca, Arthropoda, Echinodermata, and Chordata). We also observed incomplete lineage sorting in the cDNA molecular phylogeny with clades Protostomia and Deuterostomia exhibiting paraphyly (Fig. 1). However, since any bootstrap value less than 70 is considered unreliable [93,94], this node, at 17% bootstrap support, splitting the cluster taxa Chordata and Hemichordata away from Priapulida, Echinodermata, Arthropoda, Mollusca, and Brachiopoda, is not well supported. However, the node between non-bilaterians and bilaterians has 87% bootstrap support. In contrast, the amino acid molecular phylogeny (Fig. 2) shows Protostomia and Deuterostomia as monophyletic clades with 99% bootstrap support of the node at bilaterian diversification. Like the cDNA molecular phylogeny, the node at the diversification of non-bilaterians and bilaterians has 97% bootstrap support in the amino acid molecular phylogeny (Figs. 1 and 2).

**Figure 1.**
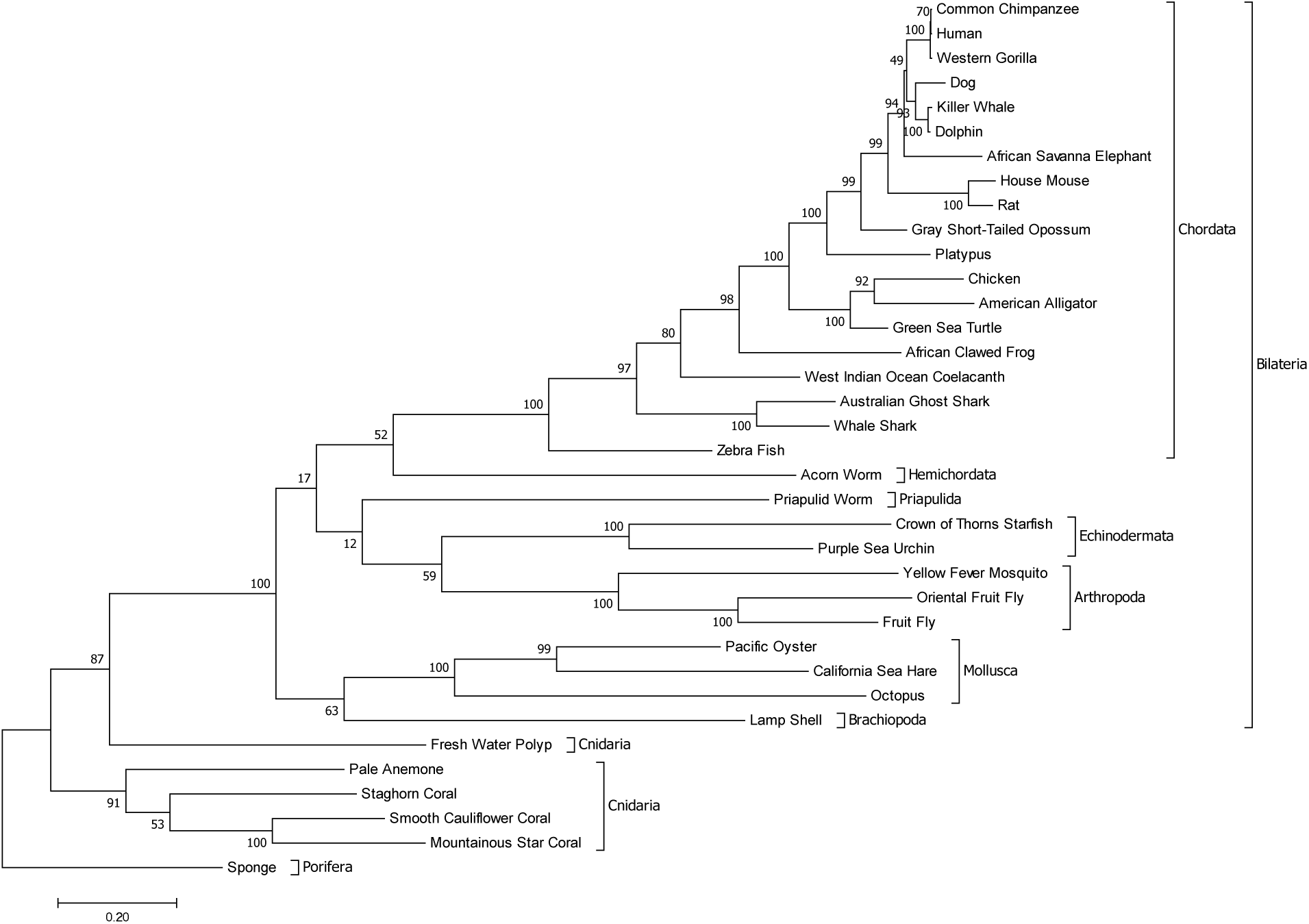
Rbm45 cDNA molecular phylogeny. Evolutionary analysis of 36 Rbm45 orthologous cDNA sequences across metazoan taxa conducted in MEGA7. The evolutionary history was inferred by using the Maximum Likelihood method based on the Tamura-Nei model [50]. The tree with the highest log likelihood (-82723.91) is shown. The bootstrap consensus tree inferred from 1000 replicates is taken to represent the evolutionary history of the taxa analyzed [51]. Branches corresponding to partitions reproduced in less than 50% bootstrap replicates are collapsed. The percentage of replicate trees in which the associated taxa clustered together in the bootstrap test (1000 replicates) are shown next to the branches. Initial tree(s) for the heuristic search were obtained automatically by applying Neighbor-Join and BioNJ algorithms to a matrix of pairwise distances estimated using the Maximum Composite Likelihood approach, and then selecting the topology with superior log likelihood value. The tree is drawn to scale, with branch lengths measured in the number of substitutions per site. There were a total of 8510 positions in the final dataset. Phyla are indicated by the inner brackets. The monophyletic clade Bilateria is indicated by the outside bracket. The tree is rooted on phylum Porifera.

**Figure 2.**
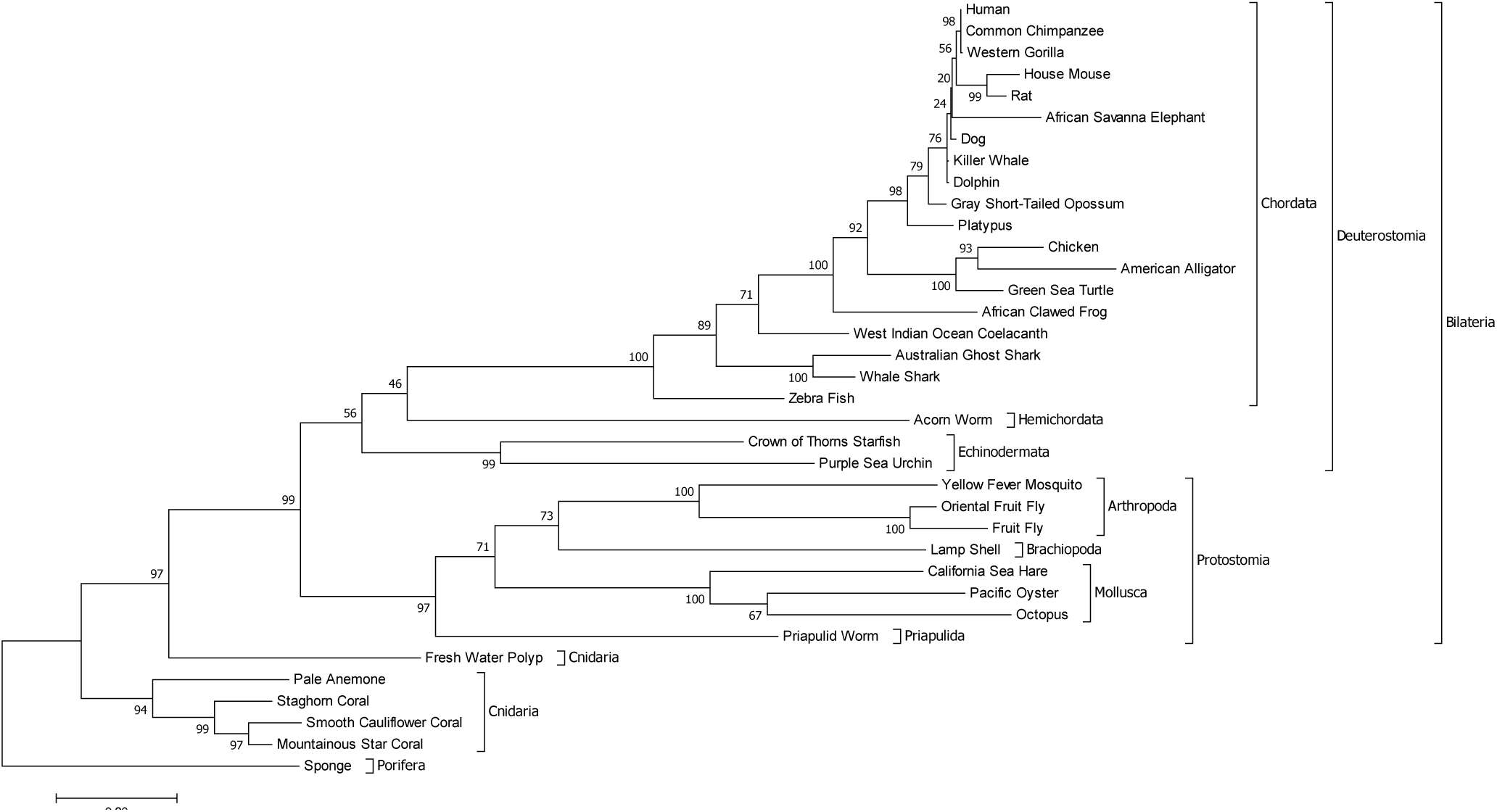
Rbm45 protein molecular phylogeny. Evolutionary analysis of 36 Rbm45 orthologous amino acid sequences across metazoan taxa conducted in MEGA7. The evolutionary history was inferred by using the Maximum Likelihood method based on the JTT matrix-based model [52]. The tree with the highest log likelihood (-22250.34) is shown. The bootstrap consensus tree inferred from 1000 replicates is taken to represent the evolutionary history of the taxa analyzed [51]. Branches corresponding to partitions reproduced in less than 50% bootstrap replicates are collapsed. The percentage of replicate trees in which the associated taxa clustered together in the bootstrap test (1000 replicates) are shown next to the branches. Initial tree(s) for the heuristic search were obtained automatically by applying Neighbor-Join and BioNJ algorithms to a matrix of pairwise distances estimated using a JTT model, and then selecting the topology with superior log likelihood value. The tree is drawn to scale, with branch lengths measured in the number of substitutions per site. There were a total of 930 positions in the final dataset. Phyla are indicated by the inner brackets. The monophyletic clades Protostomia, Deuterostomia, and Bilateria are indicated by the outside brackets. The tree is rooted on phylum Porifera.

Both the cDNA (Fig. 1) and the amino acid (Fig. 2) molecular phylogenies show phylum Hemichordata as the sister group to phylum Chordata. However, in contrast to recent work [59], neither phylogeny places Hemichordata with Echinodermata (i.e., they are paraphyletic) into clade Ambulacraria, the sister group to clade Chordata, albeit these nodes have weak bootstrap support of 52% and 46%, respectively. Conversely, in the amino acid molecular phylogeny (Fig. 2), Echinodermata and Hemichordata, though paraphyletic, do form basal groups to Chordata as expected [59,86,87]. Interestingly, both the cDNA and amino acid molecular phylogenies place Zebrafish (*Danio rerio*; group Osteichthyes; class Actinopterygii), with 100% bootstrap support, as ancestral to all other chordates (subphylum Craniata) including cartilaginous fish (class Chondrichthyes). These data are in opposition to accepted cladograms based on morphological and molecular traits [87,95], but support a hypothesis that sharks evolved from a common ancestor having a bony skeleton, with generalized bone loss as a synapomorphy for class Chondrichthyes [96]. The coelacanth (*Latimeria chalumnae*; class Sarcopterygii) has 80% (Fig. 1) and 71% (Fig. 2) bootstrap support as a sister group to clade Tetrapoda in agreement with the current hypothesis on the evolution of the tetrapod lineage [97]. Similarly, our phylogenies have 100% (Fig. 1) and 92% (Fig. 2) bootstrap support for clade Amniota.

Protostomes are separated into clades Spiralia and Ecdysozoa [86]. Within the Rbm45 amino acid molecular phylogeny (Fig. 2), which delineates clade Protostomia as monophyletic, all nodes, except for the split between octopus and Pacific oyster within phylum Mollusca, have greater than 70% bootstrap support. However, we observed incomplete lineage sorting of Rbm45 within Protostomia as clades Ecdysozoa and Spiralia do not exhibit monophyly. Instead, the taxa that make up these two clades are interspersed within the protostomes. The phylogenetic incongruences (i.e., conflicting branching orders) and lack of coalescence between our Rbm45 molecular phylogenies and currently accepted species phylogeny is not surprising [82,98] and is most likely a product of coalescent stochasticity [82,99,100] and variations in lineage-specific evolutionary rates over time (heterotachy) [101,102]. In contrast to other published data [103], we found that our amino acid molecular phylogeny recapitulated accepted species tree topology more closely than the cDNA molecular phylogeny [104]. Additionally, we have observed that where there is certainty in taxa resolution (e.g., clades Tetrapoda and Amniota), our data also appropriately resolves these nodes; whereas, where there is uncertainty in the placement of taxa (e.g., polytomy of clade Spiralia), our molecular phylogenies also reflect similar difficulties in node placement consistent with previous work by others [105].

Unfortunately, we were unable to include in our analysis two basal groups, phyla Placozoa and Ctenophora, whose positions in the metazoan phylogeny are disputed [106,107]. The predicted sequence [108] for the placozoan *Trichoplax adhaerans* Rbm45 orthologue (Gene ID: 6752484; NW_002060945.1, XM_002110678.1, XP_002110714.1; accessed 2024 February 4) is a partial sequence predicted to have at least three exons encoding 274 amino acids roughly corresponding to amino acids 21-317 (including gaps; data not shown) of human RBM45. Therefore, the hypothetical *T. adhaerans* Rbm45 orthologue sequence lacks a start codon, RNA-binding domain III, homo-oligomer assembly domain, nuclear localization signal, and a stop codon. However, the presence of at least 3 exons supports the assertion that placozoans are more derived than their simple body plan would imply [106,108] (Fig. 3). Inclusion of the *T. adhaerans* partial Rbm45 amino acid sequence into our molecular phylogenetic analysis placed it as a sister group to phyla Porifera and Cnidaria as expected [86,87] (data not shown). The nuclear genomes of six species from phylum Ctenophora have been sequenced to date (https://www.ncbi.nlm.nih.gov/search/all/?term=Ctenophora; accessed 2024 February 4); however, an Rbm45 orthologue has not yet been identified. This is unfortunate, as we wished to include this gene product in our analysis considering the hypothesis [107,109] that ctenophores are the sister group to all animals, and not poriferans, challenging the predominant paradigm that metazoan nervous systems have evolved complexity in a stepwise manner over deep time [110].

**Figure 3.**
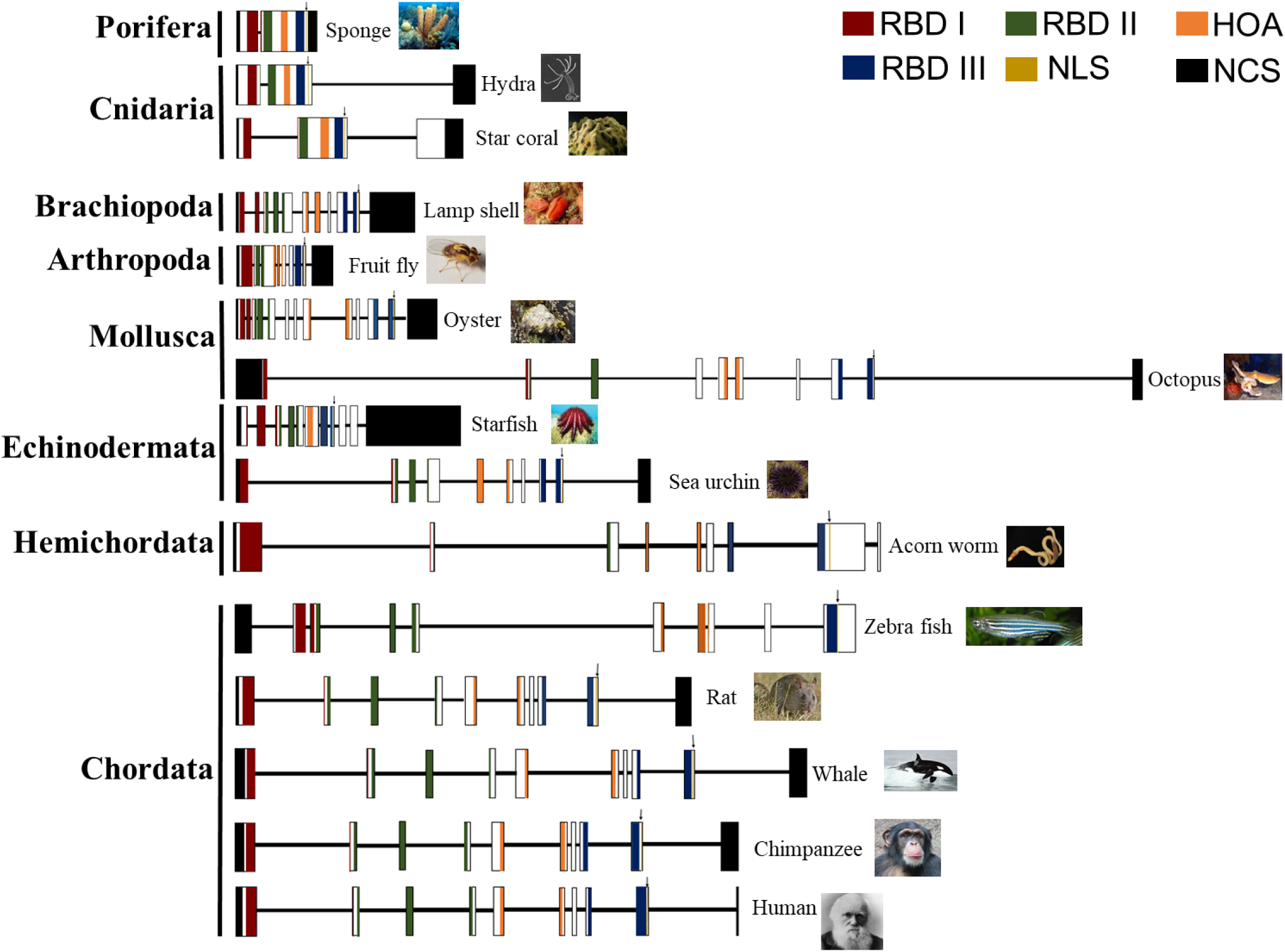
Gene architecture of Rbm45 orthologues. Rbm45 protein domains are linearly conserved from sponges to humans. Schematic diagram of the exon-intron structure from 15 representative Rbm45 orthologues across metazoan taxa. Vertical boxes represent exons, and the solid horizontal lines represent introns. The diagram shows the relative sizes of the exons and introns. The introns and exons are scaled by type; the intron width scale is half that of the exon width scale. The DNA sequences encoding RNA-binding domains (RBD) I, II, and III, and the homo-oligomer assembly (HOA) domain, are indicated by red, green, blue, and orange fill, respectively. The DNA sequences encoding the monopartite nuclear localization signal (NLS) are indicated by a vertical yellow line and a downward pointing arrow. Non-coding exon sequences (NCS) are shaded black. All animal images were retrieved from Wikimedia (commons.wikimedia.org) and are public domain (Creative Commons license CC0: https://creativecommons.org/).

### Rbm45 Orthologue Protein Domain Conservation

Considering the strong recapitulation of the metazoan tree of life using Rbm45 amino acid sequences, we also analyzed the conservation of protein domains from sponges to humans across the same 36 taxa used in the phylogenetic analysis (Figs. 1 and 2). Multiple sequence alignment using the Clustal Omega algorithm demonstrates that Rbm45 protein domains are linearly conserved from sponges to humans in the order: RBD I, RBD II, HOA, RBD III, and NLS (Fig. 3). RBDs I, II, and III show 72%, 80%, and 65 % similarity across taxa, respectively, with the HOA domain displaying 68% similarity across taxa analyzed (data not shown). Furthermore, the monopartite NLS [32,111], located just downstream of RBD III (Fig. 3), from all 36 orthologues conforms to the canonical consensus sequence K(R/K)(X)(R/K). Example NLSs are sponge KRQK, sea anemone KRPR, mosquito KRMR, and human KRQR (data not shown) exemplifying core basic amino acids.

In contrast to the NLS, the human RBM45 NES does not conform to a classical leucine-rich domain but instead was empirically determined [32] to be made up of a clique of two hydrophobic amino acids: leucine-leucine. Additionally, this clique of two hydrophobic residues L(L/I) are conserved among Rbm45 mammalian orthologues [27,32]. Using multiple sequence alignment (Materials and Methods), we identified the L(L/I) clique only within clade Tetrapoda of clade Craniata. To identify an NES in non-tetrapods, we interrogated the online NES prediction program LocNES [56] using human RBM45 as the input sequence. The algorithm identified a majority-rule canonical sequence from all 19 Craniata taxa (i.e., clade Tetrapoda, clade Chondrichthyes, and group Osteichthyes) between the HOA and RBD III domains [31,32] having the general form: R^15^K^16^MA(T^14^/S^2^)Q(M^12^/L^7^)VA^16^A^18^Q^18^(L^11^/M^5^/V^3^)(A^11^/M^4^)(S^18^/T^1^)(M^15^/V^3^) where the superscript indicates the number of taxa with that amino acid, if not at identity, with conserved amino acids (e.g., hydrophobic) grouped in parentheses. In tetrapods, this conserved sequence was downstream and immediately adjacent to the L(L/I) clique. In contrast, using sponge or sea anemone Rbm45 as the query sequence, we were unable to deduce a consensus NES in invertebrates (clades Ambulacraria and Protostomia). These data agree with a pairwise alignment of sponge and human Rbm45 amino acid sequences where the Clustal Omega algorithm inserted a gap in the sponge sequence across from the human NES (data not shown). The inability to identify a putative NES in the invertebrate taxa analyzed is not necessarily surprising given the complexity and variability of NES sequences [56,112]. Furthermore, the LocNES algorithm searched for CRM1 dependent NESs [56]. It is possible that invertebrate Rbm45 proteins use a CRM1-independent pathway (e.g., passive diffusion) like vertebrate TDP-43 and FUS [113]. If invertebrate Rbm45 exits the nucleus by passive diffusion, then the evolution of an NES would be a novel trait in the Craniata lineage. Future work on this question would necessitate the empirical determination of whether Rbm45 is able to be trafficked from the nucleus to the cytoplasm in these invertebrate organisms.

### Rbm45 Orthologue Gene Architecture Evolution

Concurrent with protein domain conservation analysis, we also analyzed the gene structure of 25 *Rbm45* orthologues (Materials and Methods). *Rbm45* from the non-bilaterian phylum Porifera has 2 large exons, and from phylum Cnidaria hydra and star coral have 3 exons (Fig. 3) and the pale anemone has 4 exons, including a cryptic exon (data not shown).

Interestingly, all protein domains are conserved in the first two exons of cnidarian *Rbm45* with a similar spacing to sponge (Fig. 3 and data not shown). The bilaterian phyla have between 6 and 13 exons, with the Rbm45 protein domains fragmented across multiple exons (Fig. 3 and data not shown). Unsurprisingly, though the localization of protein domains occurs in different exons depending on the organism, within each phylum the domains are localized in very similar exon positions. This phenomenon is especially evident in class Mammalia of phylum Chordata where the domain distribution among exons is almost identical (Fig. 3 and data not shown) between taxa. Furthermore, regression analysis of the 25 *Rbm45* orthologues revealed a statistically significant strong negative correlation between mean exon length and total number of exons (R^2^ = 0.6169, *P* < 0.0001; Fig. 4) in accord with the work of others [114]. Our data are in agreement with earlier work demonstrating that more evolutionarily advanced organisms, as measured by genomic and metabolomic complexity [114–117], have more short exons and longer introns while less evolutionary advanced organisms have fewer large exons and short introns (Fig. 3 and data not shown) consistent with whole genome analysis across the five kingdoms [118] Protozoa, Chromista, Plantae, Fungi, and Animalia [114].

**Figure 4.**
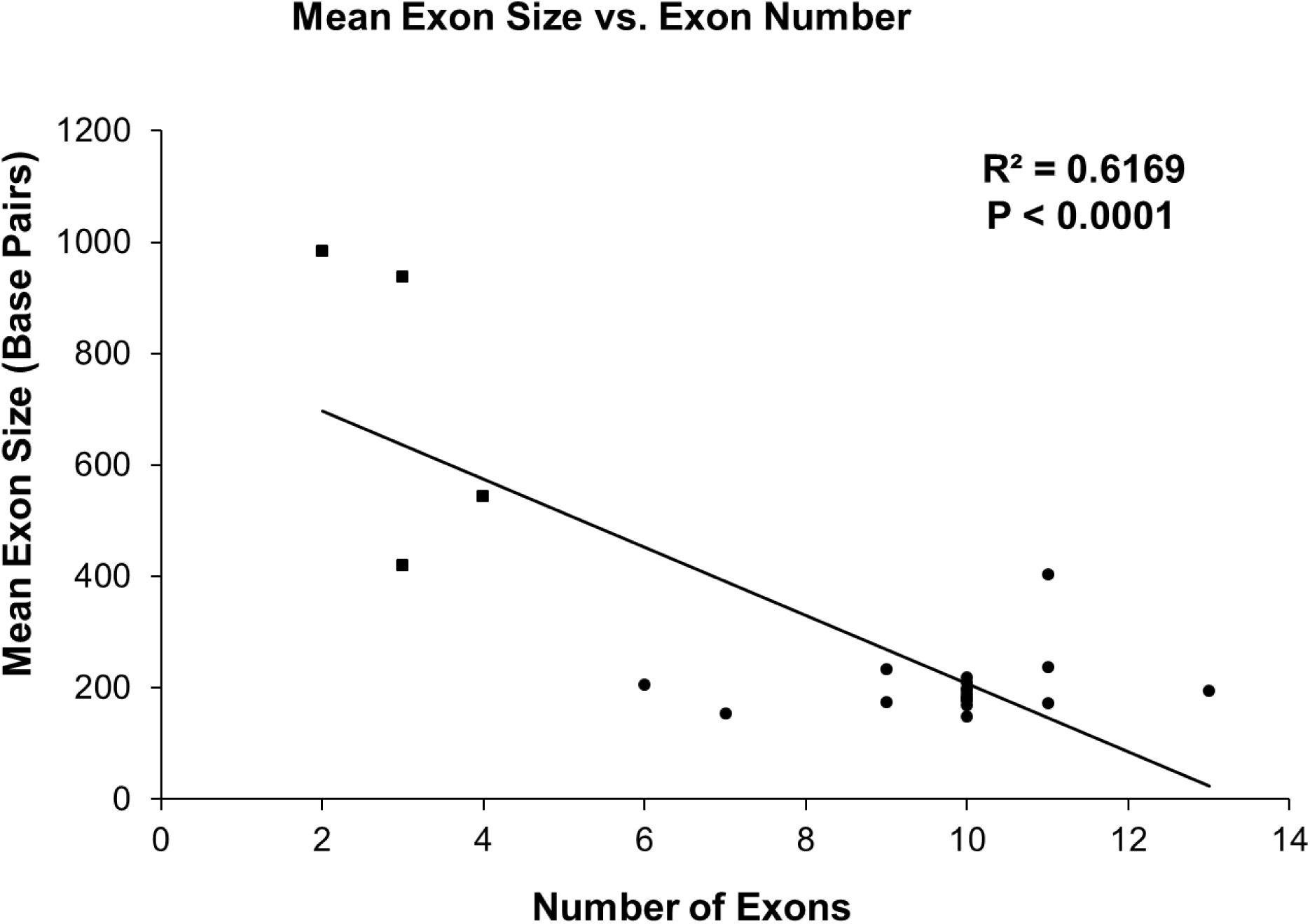
Regression analysis of mean exon length versus number of exons in Rbm45 orthologues. Rbm45 mean exon length decreases as exon number increases. A scatter plot, with linear regression, was produced from 25 representative Rbm45 orthologues. There is a statistically significant correlation (Coefficient of Determination: R^2^) between mean exon length and the number of exons in the organism’s gene structure. Closed squares (▪) are non-bilaterian (i.e., diploblastic) animals of phyla Porifera and Cnidaria; closed circles (•) are members of clade Bilateria (i.e., triploblastic) from phyla Arthropoda, Brachiopoda, Chordata, Echinodermata, Hemichordata, and Mollusca [86].

We extended our analysis of the evolution of *Rbm45* gene complexity by examining the correlation of a taxon’s approximate lineage age, as determined by a robust analysis of current literature (Materials and Methods), to the number of exons in the *Rbm45* orthologue. We demonstrate a statistically significant very strong negative correlation (R^2^ = 0.8057, *P* < 0.0001; Fig. 5) where the most ancient lineages (e.g., Porifera at 650,000,000 years; Cnidaria at 500-570,000,000 years; see Materials and Methods) have the fewest number of exons, between 2 and 4, while generally more recent taxa lineages have more exons (> 9 exons; e.g., Zebrafish at 150,000,000 years; crown-of-thorns starfish at 4,000,000 years; humans at 300,000 years; see Materials and Methods). The majority of taxa followed the trend of fewer exons correlating to an ancient lineage age and more exons to a more recent lineage age. A notable exception is the acorn worm, *Saccoglossus kowalevskii*, from phylum Hemichordata (clade Ambulacraria) which has an approximate lineage age of 370,000,000 years (Upper Devonian) [59]. The reference genomic sequence (NW_003156738.1) of *S. kowalevskii* predicts 9 exons which is closer in number to what is observed for other members of clade Bilateria we analyzed. However, all other bilaterian taxa that we used in Fig. 5 had lineage ages of less than 250,000,000 years (e.g., horseshoe crab with 6 exons). A rigorous analysis of the gene structure of all 36 *Rbm45* orthologues used in this study revealed that the *Priapulis caudatus* (phylum Priapulida; clade Scalidophora) *Rbm45* orthologue reference genomic sequence (NW_014578398.1) is predicted to have 17 exons (data not shown), the most of any organism examined by us. Extant priapulins date from the late Carboniferous (∼350,000,000 years ago) [119], while extinct stem- and crown-group priapulins are found in the middle Cambrian (∼500,000,000 years ago) [120,121]. These data suggest that division of the *Rbm45* gene into many exons (i.e., > 4) occurred relatively early in the adaptive radiation of the evolutionary complex bilateral body plan during and after the Cambrian period [122,123]. Taken together, these data indicate an ancient origin for *Rbm45* in the metazoan lineage. In accordance with this observation, we were able to identify an *Rbm45* orthologue in *Monosiga brevicollis* [(Monbr1│Name: e_gw1.8.135.1; Protein ID: 16822; Location: scaffold_8:72832-74415); https://mycocosm.jgi.doe.gov/cgi-bin/dispGeneModel?db=Monbr1&id=16822] from phylum Choanoflagellata, clade Holozoa. These flagellated protists are hypothesized to be the ancestors of phyla Porifera, and thus all metazoans [124–126], further demonstrating the ancient roots of *Rbm45*. Consistent with this well-accepted hypothesis [86], an unrooted phylogeny using amino acid sequence data from the 36 Rbm45 orthologues from this study plus the Rbm45 orthologue from Choanoflagellata places choanoflagellates at the root of the tree as the sister to all animals (data not shown). This high level of conservation among crown clades reveals that *Rbm45* may play a role in neurogenesis across metazoans.

**Figure 5.**
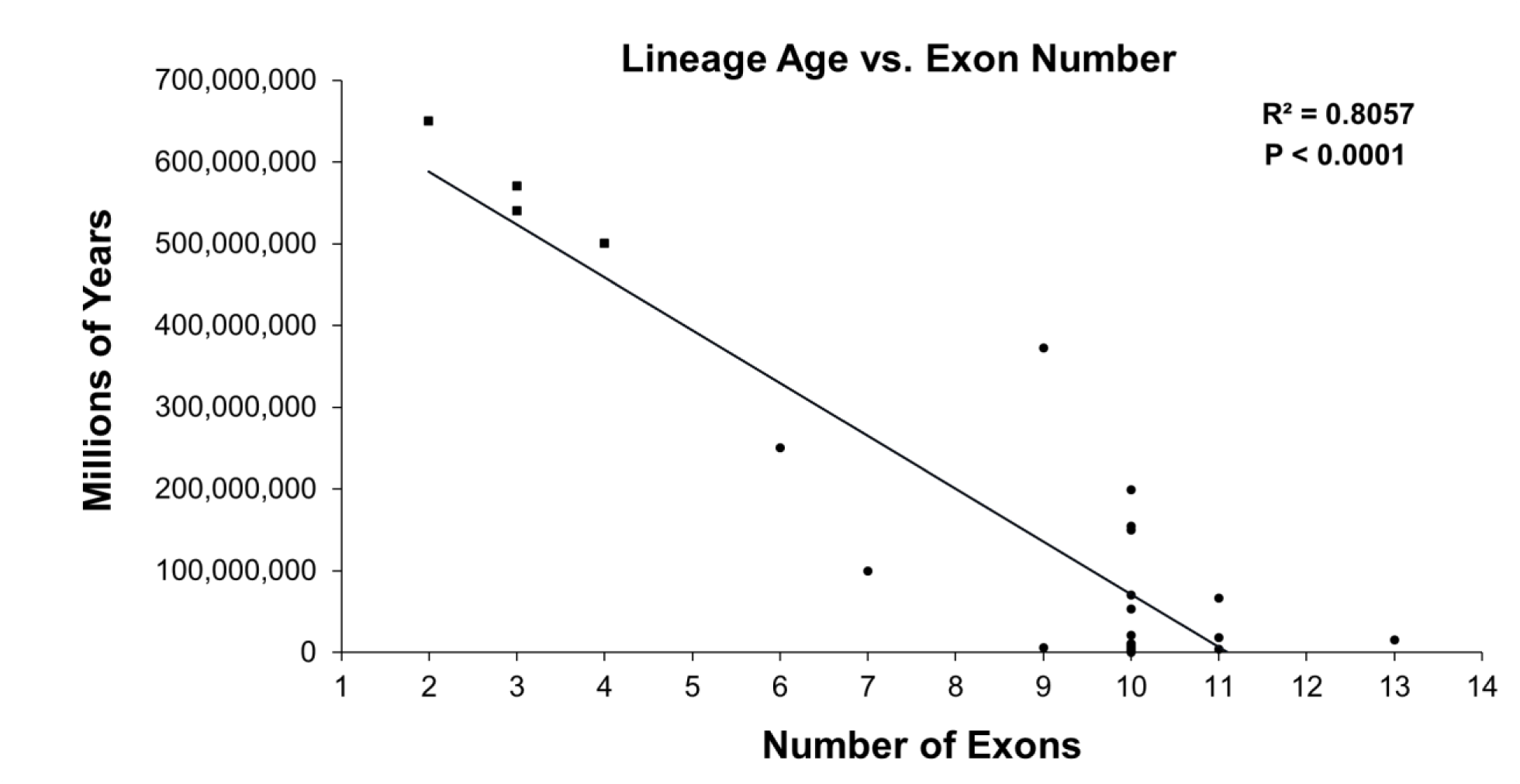
Regression analysis of species lineage age versus number of exons in Rbm45 orthologues. Rbm45 exon number increases as lineage age decreases. A scatter plot, with linear regression, was produced from 25 representative Rbm45 orthologues. There is a statistically significant high correlation (Coefficient of Determination: R^2^ ≥ 0.7) between lineage age and the number of exons in the organism’s gene structure. Closed squares (▪) are non-bilaterian (i.e., diploblastic) animals of phyla Porifera and Cnidaria; closed circles (•) are members of clade Bilateria (i.e., triploblastic) from phyla Arthropoda, Brachiopoda, Chordata, Echinodermata, Hemichordata, and Mollusca [86].

## Conclusions

We have demonstrated that *Rbm45* is an ancient gene conserved from clade Holozoa to clade Metazoa with phylogenetic analysis of Rbm45 orthologue amino acid sequence mirroring known monophyletic relationships among metazoans. Additionally, we have verified and extended the observation of deep homology of the RBD, HOA, and NLS regions in the Rbm45 protein from sponges to chordates, whereas the NES is a possible synapomorphy unique to clade Craniata. Furthermore, we have shown a statistically significant increase in complexity of *Rbm45* gene architecture contemporaneous with increasing evolutionary complexity moving from non-bilaterian to bilaterian animals over evolutionary time. Elucidation of Rbm45 function in neural development and homeostasis in a broad range of molecular genetic model systems/taxa will merit detailed attention in the future to holistically understand its function in a breadth of neural/sensory networks.

## *Abbreviations

Rbm45: RNA-binding motif protein 45;
RBP: RNA-binding protein;
RRM: RNA recognition motif;
RBDP: RNA recognition motif-type binding domain protein;
RBD: RNA-binding domain;
HOA: homo-oligomer assembly;
NLS: nuclear localization signal;
NES: nuclear export signal;
NCBI: National Center for Biotechnology Information; no., number;
MEGA7: Molecular Evolutionary Genetic Analysis v.7.0 software;
JTT: Jones-Taylor-Thornton model;
LocNES: Locating Nuclear Export Signals algorithm;
IUCN: International Union for Conservation of Nature;
NCS: non-coding exon sequence

## Acknowledgements

We thank Trevor Butler for helpful discussions on animal lineage age and exon number. J.O.H thanks Julie K. Henderson for editorial assistance. This work was supported by funds from the Judson University science department (V.V., T.N.M., A.C., L.M.S., J.O.H.), the William W. Brady Chair of Science endowment (J.O.H), and two one-semester sabbatical leaves provided by Judson University (J.O.H: 2017 and 2024).

## Author contributions

Conceptualization: J.O.H.; Formal Analysis: V.V., A.C., and J.O.H.; Investigation: V.V., T.N.M., A.C., L.M.S., and J.O.H.; Supervision: J.O.H.; Visualization: V.V., T.N.M., A.C., and J.O.H.; Writing – original draft: J.O.H.; Writing – reviewing and editing: V.V., T.N.M., A.C., and J.O.H.

